# Neural circuit mechanism underlying the feeding controlled by insula-central amygdala pathway

**DOI:** 10.1101/750562

**Authors:** Calvin Zhang-Molina, Matthew B Schmit, Haijiang Cai

## Abstract

Central nucleus of amygdala (CeA) contains distinct populations of neurons that play opposing roles in feeding. The circuit mechanism of how CeA neurons process information sent from their upstream inputs to regulate feeding is still unclear. Here we show that activation of the neural pathway projecting from insular cortex neurons to CeA suppresses food intake. Surprisingly, we find that the inputs from insular cortex form excitatory connections with similar strength to all types of CeA neurons. To reconcile this puzzling result, and previous findings, we developed a conductance-based dynamical systems model for the CeA neuronal network. Computer simulations showed that both the intrinsic electrophysiological properties of individual CeA neurons and the overall synaptic organization of the CeA circuit play a functionally significant role in shaping the CeA neural dynamics. We successfully identified a specific CeA circuit structure that reproduces the desired circuit output consistent with existing experimentally observed feeding behaviors.

## Introduction

Neurons in the central nucleus of the amygdala (CeA) play an important role in controlling feeding, a behavior critical to our survival and health (Cai et al., 2014; Douglass et al., 2017; Hardaway et al., 2019; Ip et al., 2019; Petrovich et al., 2009). However, how the CeA neurons control food intake at circuit level is still poorly understood. CeA is primarily composed of γ-aminobutyric acid (GABAergic) inhibitory neurons, which can be classified into multiple different types based on their gene expression profiles or distinct electrophysiological properties (Chieng et al., 2006; Ciocchi et al., 2010; Day et al., 1999; Fadok et al., 2018; Haubensak et al., 2010; Kim et al., 2017; Lopez de Armentia and Sah, 2004; McDonald and Augustine, 1993; Sah et al., 2003). Using genetic markers, several recent studies identified at least two distinct populations of neurons in CeA that play opposing roles in controlling feeding (Cai et al., 2014; Douglass et al., 2017). Specifically, one study found that activation of the neurons marked by the expression of protein kinase C-delta (PKC-δ+) suppresses food intake while silencing these neurons increases food intake in fed state (Cai et al., 2014). Another study identified a different population of neurons marked by the expression of the serotonin receptor 2a (Htr2a+), which do not overlap with PKC-δ+ neurons (Douglass et al., 2017). In contrast to PKC-δ+ neurons, activation of the Htr2a+ neurons increases food intake while silencing these neurons suppresses food intake (Douglass et al., 2017). Particular physiological properties give rise to distinctive discharge patterns and might play important roles in controlling functions of the neurons during behaviors. Based on their action potential firing in response to injection of depolarization currents, neurons in CeA are usually classified into three types: 1) late firing neurons and 2) regular spiking neurons which are mostly observed in the lateral part of CeA, and 3) low threshold bursting neurons which are usually in the medial part of CeA (Chieng et al., 2006; Dumont et al., 2002; Lopez de Armentia and Sah, 2004). Interestingly, both PKC-δ+ and Htr2a+ neurons are primarily located in the lateral part of CeA and contain mostly late firing neurons and a small proportion of regular spiking populations (Cai et al., 2014; Douglass et al., 2017). Thus, the distinct functions in feeding of these two populations of neurons cannot be explained by their electrophysiological properties. Another possibility for these neurons to have distinct functions is that they receive inputs from different upstream brain regions, and thereby activated in different situations to control feeding. One dominant top-down cortical input to CeA and other amygdala subnuclei is from the insula (Fudge and Tucker, 2009; McDonald et al., 1999; Mufson et al., 1981; Shi and Cassell, 1998), neurons in which play important roles in processing taste and visceral information (Accolla and Carleton, 2008; Andermann and Lowell, 2017; Augustine, 1996; Caruana et al., 2011; Chen et al., 2011; Craig, 2003; Gogolla, 2017; Katz et al., 2001; Samuelsen and Fontanini, 2017; Yamamoto et al., 1985). Recent studies reported that the projection from insular cortex to CeA signals aversive bitter taste as tested in drinking behaviors (Schiff et al., 2018; Wang et al., 2018), suggesting this pathway might also control feeding behavior. However, whether the insula-CeA pathway regulates solid food intake and what type of CeA neurons are innervated by the inputs from insular cortex are still unknown.

Here we find that optogenetic activation of the insula-CeA pathway strongly and rapidly suppresses feeding. Surprisingly, our optogenetics-assisted circuits mapping show that insular cortex neurons send monosynaptic excitatory inputs non-selectively to all CeA neurons with similar strength, independent of their genetic markers or electrophysiological properties. Based on these new results and previously published data of CeA neurons, we developed a conductance-based dynamical systems model for the CeA circuit to explore the possible underlying circuit structure of CeA neurons in feeding control. Interestingly, out of the ten fundamentally different circuit structures considered, only one specific circuit structure was able to consistently generate the observed feeding behaviors. Computer simulations of our CeA circuit model revealed that both the regular spiking and the late firing CeA neurons play an essential role in modulating the CeA circuit function, and only one specific combination of the individual CeA neurons’ electrophysiological properties and the CeA circuit’s synaptic organization can reproduce the desired circuit output.

## Results

### Activation of the projection from insular cortex to CeA suppresses feeding

In order to specifically target the insular cortex neurons that project to the CeA, we used a two-virus Cre-on strategy, in which we stereotaxically injected Cre-dependent adeno-associated virus (AAV) encoding channelrhodopsin (ChR2) (Zhang et al., 2007) or enhanced yellow fluorescent protein (EYFP) control into the insular cortex and AAVretro-Cre (Tervo et al., 2016) into the CeA bilaterally in wild type mice. Meanwhile, we implanted optic fibers above the CeA. Thus, only the nerve terminals in CeA that projected from insular cortex neurons express ChR2 and can be activated by light stimulation (Fig. 1A). Histological analysis revealed that ChR2-expressing neurons were restricted to a sub-region of the insular cortex (Fig. 1B, C). Interestingly, the fluorescent nerve terminals projected from these insular cortex neurons were remarkably exclusive to the lateral part of CeA but not to the surrounding regions (Fig. 1D). 3-4 week after virus expression and animal recovery, we coupled the optic fibers to a blue laser (473 nm) to stimulate the CeA terminals projected from insular cortex neurons. We found that photo-stimulation of these terminals reduced total amount of food intake in both fed (Fig. 1E) and 24-hr fasted animals (Fig. 1F). Light activation of this pathway also significantly increased the latency to eat (Fig. S1), suggesting the initiation of feeding is also inhibited. Light stimulation of the insula-CeA neural pathway does not affect the movement or anxiety levels significantly as tested in an open field assay (Fig. 1I-L). We also tested feeding when the mice were in their home cages, where the mice were less anxious. Mice were 24-hr fasted and either blue light or control light (593 nm), which does not activate ChR2, was delivered one to two seconds after the onset of eating. We found that light stimulation of the insula-CeA pathway significantly shortened the duration of feeding bouts (Fig. 1G, H), indicating that feeding was suppressed in the home cage. These results suggest that insular cortex stimulation is able to reduce total food intake as well as bout duration, and that this is not a result of anxiety or of any impaired mobility.

**Fig. 1.**
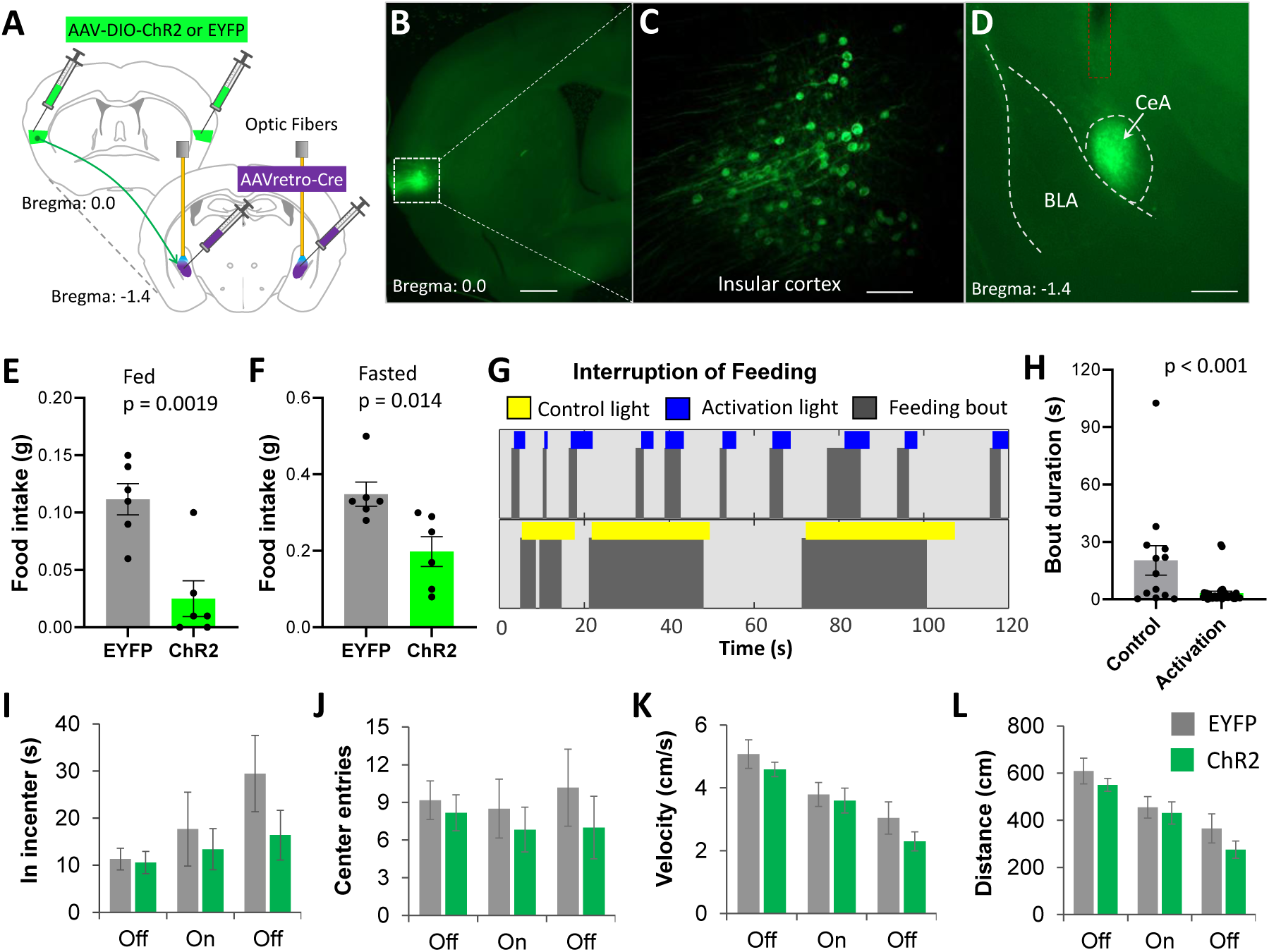
Optogenetic activation of the insula-CeA projection suppresses food intake. A. Diagram shows virus injection strategy and optic fiber implantation. B. ChR2-EYFP expression in insular cortex neurons verified in histology post experiments. Bar, 500 µm C. High resolution image show EYFP expression in individual insular cortex neurons. Bar, 100 µm. D. EYFP fluorescent nerve terminals are restricted in CeA region. Red dotted line outlines the optic fiber tract. Bar, 500 µm. E. Food intake in fed mice after light stimulation of the insula-CeA projections in a feeding session of 30 min. Light pulse 10 ms, 10 Hz. Unpaired *t*-test, t_(10)_ = 4.19. n = 6 animals in each group. F. Food intake in 24-hr fasted mice after light stimulation of the insula-CeA projections in a feeding session of 20 min. Light pulse 10 ms, 10 Hz. Unpaired *t*-test, t_(10)_ = 2.99. n = 6 animals in each group. G. A representative raster lot show that feeding bouts are interrupted by light stimulation. Light pulses were usually delivered 1-2 seconds after the feeding was started. Mice were 24 hr fasted and tested in their home cages. H. Quantification of the feeding bout duration after light stimulation. Unpaired *t*-test, t_(48)_ = 3.56. n = 13 bouts in control and 37 bouts in activation group. I-L. Light stimulation does not impair movement or anxiety in the open field test. Light pulses (10 ms, 10 Hz, 2 min) were delivered after 2 min no light base line and followed by 2 min post light. Two-way ANOVA. No difference between EFYP and ChR2 detected. n = 6 animals in each group. Data shown as mean ± s.e.m.

### CeA neurons receive monosynaptic excitatory inputs from insular cortex

To determine if CeA neurons are monosynaptically innervated by inputs from insular cortex neurons, we stereotaxically injected and AAV encoding ChR2-EYFP in insular cortex in wild type mice. After allowing 3-4 weeks for post-surgical recovery and virus expression, we prepared live brain slices and performed whole-cell patch clamp recording on neurons in the CeA (Fig. 2A-C). The brain sections that contained insular cortex were also cut to verify the ChR2-EYFP expression in insular cortex neurons. We found that the EYFP positive nerve fibers were restricted to the CeA region (Fig. 2B). Although the soma of the neural projections from insular cortex were cut away in the brain slices with the CeA, we found that optogenetic activation of the ChR2-expressing terminals in CeA is sufficient to trigger postsynaptic responses in CeA neurons. When the cells are voltage-clamped at −70 mV, we observed robust light-triggered excitatory postsynaptic currents (EPSCs), which can be blocked by the competitive AMPA/kainate receptor antagonist CNQX (Fig. 2D). All these EPSCs are triggered within a short delay of less than 4 ms after the light pulses (Fig. 2D inset), suggesting the connection is monosynaptic. Almost all the CeA neurons tested (32 out of 33 tested cells) show EPSCs in response to light pulses. This result is consistent with a previous study that reported nearly all the CeA neurons receive excitatory innervation from insular cortex (Schiff et al., 2018), and also consistent with the previous monosynaptic retrograde rabies tracing studies, which demonstrated that both CeA PKC-δ+ neurons and CeA Htr2a+ neurons receive monosynaptic inputs from neurons in insular cortex (Cai et al., 2014; Douglass et al., 2017). Interestingly, when we voltage-clamped the CeA neurons at −40 mV, we observed robust upward inhibitory postsynaptic currents (IPSCs) following the downward EPSCs, both of which are blocked by CNQX (Fig. 2E). Given the fact that all CeA neurons are GABAergic inhibitory neurons (Haubensak et al., 2010; Sun and Cassell, 1993) and the mutual inhibition between CeA neurons described in previous studies (Hou et al., 2016; Hunt et al., 2017), these results suggest that the IPSCs are disynaptic to the insular cortex inputs and are the result of GABAergic inhibition from other CeA neurons activated by light-stimulated insular cortex inputs (Fig. 2F).

**Fig. 2.**
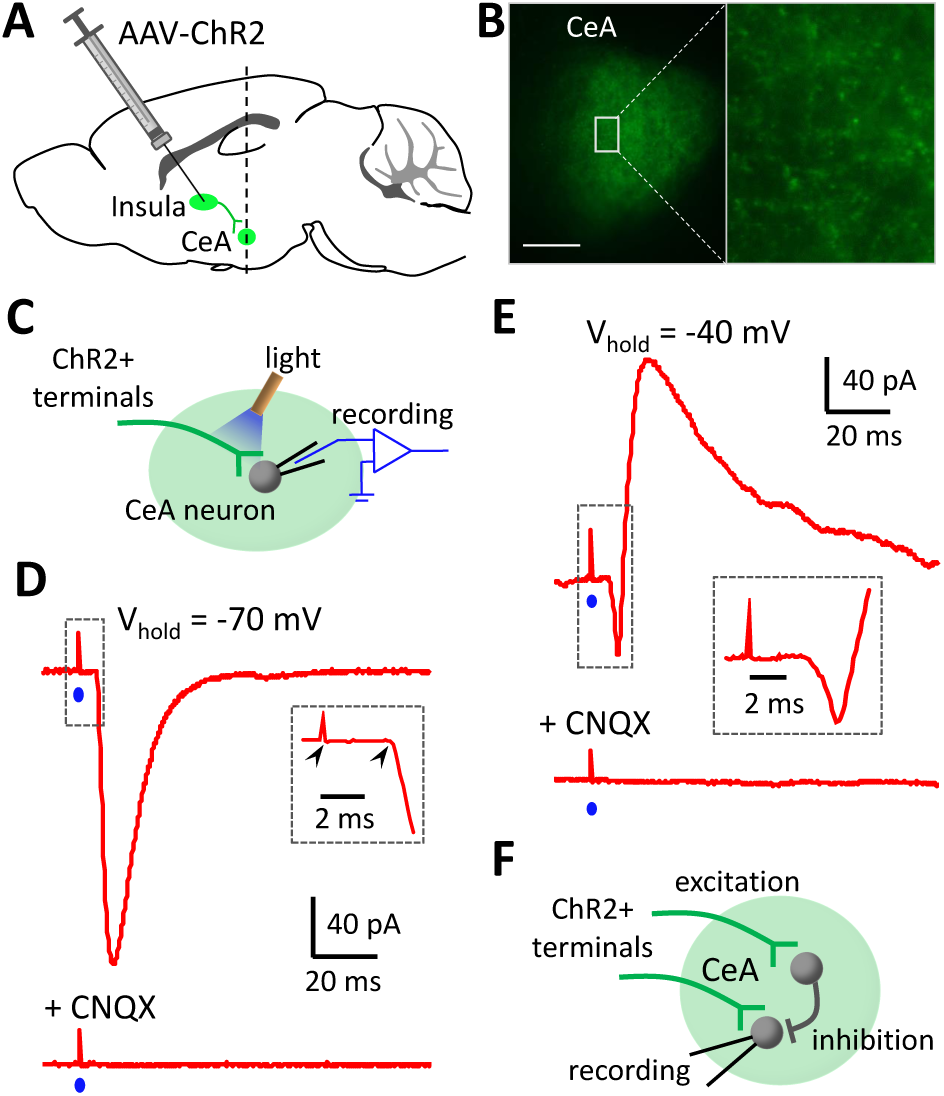
CeA neurons receive monosynaptic excitatory inputs from insular cortex. A. Diagram shows virus injection and brain slice with CeA were cut for electrophysiological recording. B. A representative image show that the fluorescent nerve terminals are located in CeA of live brain slices. Bar, 200 µm. C. Diagram shows light stimulation of the ChR2-expressing nerve terminals and electrophysiological recording on CeA neurons. D. Sample recording traces show that light pulse (blue dot) triggers EPSC in a CeA neuron when it is voltage-clamped at −70 mV and this EPSC is blocked by bath application of 20 µM CNQX. Inset, the latency of the EPSC is measured from the start of light to the start of inward current (arrows). E. Sample recording traces show that light pulse triggers both EPSC and IPSC in a CeA neuron when it is voltage-clamped at −40 mV. Both the EPSC and IPSC are blocked by CNQX (20 µM). F. The circuit connection suggested by the electrophysiological recording results.

### Insular cortex neurons innervate CeA neurons non-selectively

Because activation of the insular-CeA pathway suppresses food intake, an effect the same as activation of CeA PKC-δ+ neurons, we hypothesize that the excitation from insula cortex to CeA PKC-δ+ neurons (PKC-δ+) is stronger than that in PKC-δ negative (PKC-δ-) neurons. In order to identify the PKC-δ+ neurons during slice recording, we crossed the PKC-δ-Cre mice with the AI14 Cre-reporter line (Madisen et al., 2010) to express tdTomato in PKC-δ+ neurons. Our previous study has shown that almost all the CeA PKC-δ+ neurons (>99%) are labelled by tdTomato in this way (Cai et al., 2014). With this strategy, we can identify CeA PKC-δ+ neurons by their tdTomato expression in live brain slices (Fig. 3A). We stereotaxically injected an AAV encoding ChR2-EYFP in the insular cortex of the PKC-δ-Cre-AI14 crossed mice. 3-4 weeks after virus injection, we performed slice electrophysiological recording on CeA neurons. We first current-clamped the neurons and applied a series of depolarization steps to elicit action potential firing, which were used to characterize electrophysiological properties of the recorded neurons (Fig. 3B-D). We then voltage-clamped the neurons and elicited EPSCs by photo-stimulating the nerve terminals projected from insular cortex. Although there is a trend towards larger EPSCs in PKC-δ+ neurons than in PKC-δ-neurons, we did not observe a significant difference (Fig. 3E). The EPSC latency in both populations was around 3 ms (Fig. 3F), suggesting monosynaptic innervation. We also checked the electrophysiological properties of these neurons (Fig. 3G-I). Our only observed difference was a significantly longer delay in action potential firing in response to the injection of a depolarization current (Fig. 3B, G), which is consistent with a previous report that shows majority of the PKC-δ+ neurons are late firing neurons (Haubensak et al., 2010).

**Fig. 3.**
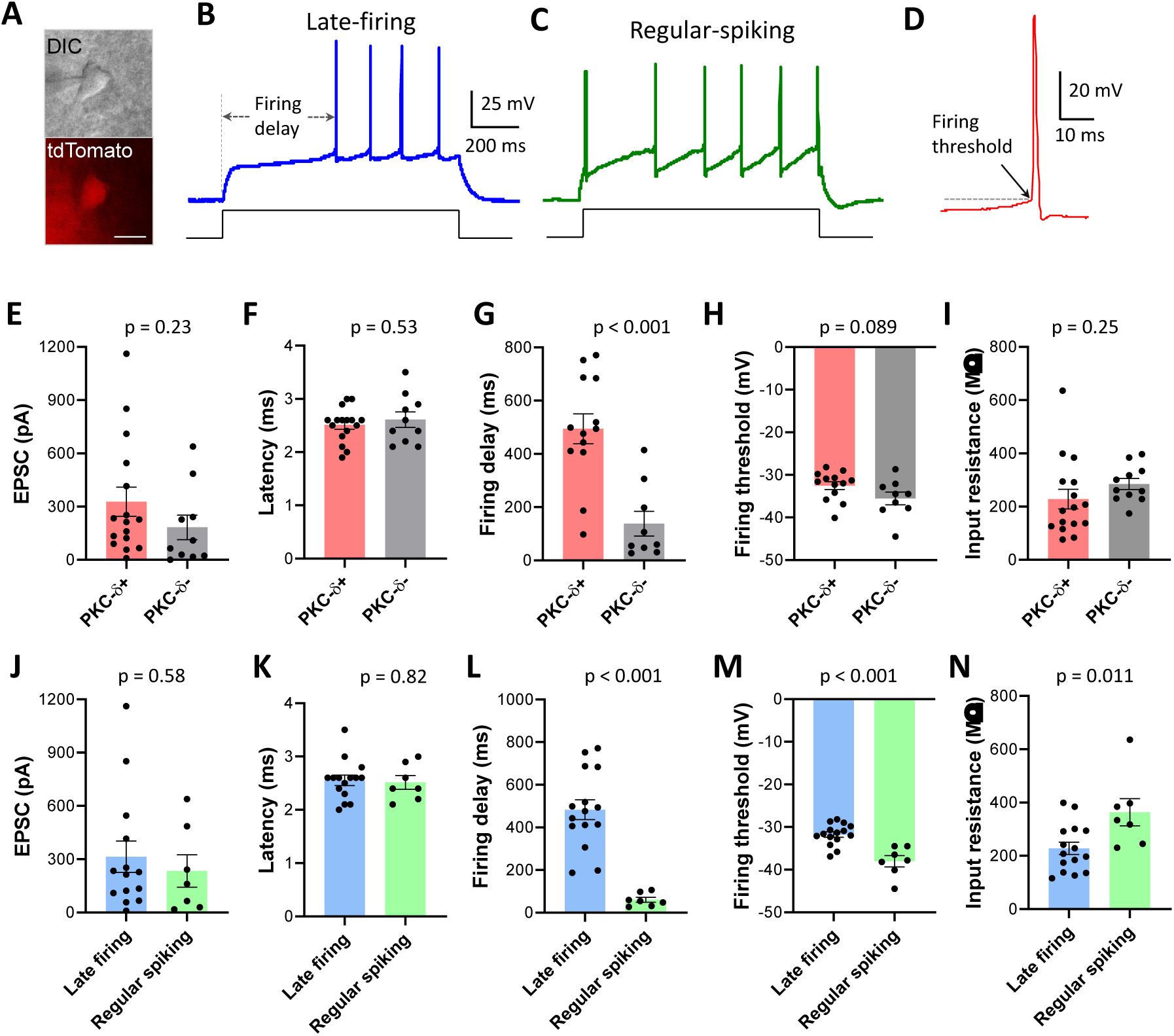
Insular cortex innervates CeA neurons nonselectively. A. PKC-δ+ neurons can be identified by tdTomato expression in brain slices. DIC, differential interference contrast. Bar, 10 µm. B. Sample recording trace of a late-firing neuron in response to a step of depolarizing current injection. C. Sample recording trace of a regular spiking neurons in response to a step of depolarizing current injection. D. Firing threshold identified in a sample recording trace of an action potential. E-I. EPSC (E), EPSC latency (F), action potential firing delay (G), action potential firing threshold (H) and input resistance (I) in PKC-δ+ neurons and PKC-δ-neurons. Unpaired *t*-test. t_(24)_ = 1.23 (E), t_(24)_ = 0.64 (F), t_(20)_ = 4.59 (G), t_(20)_ = 1.79 (H), t_(25)_ = 1.18 (I). n = 13-16 PKC-δ+ neurons and 9-11 PKC-δ-neurons. J-N. EPSC (J), EPSC latency (K), action potential firing delay (L), action potential firing threshold (M) and input resistance (N) in late-firing neurons and regular spiking neurons. Unpaired *t*-test, t_(19)_ = 0.57 (J), t_(20)_ = 0.23 (K), t_(20)_ = 6.07 (L), t_(20)_ = 4.84 (M), t_(20)_ = 2.80 (N). n = 14-15 late-firing neurons and 7 regular-spiking neurons. Data shown as mean ± s.e.m.

Neurons in the CeA are usually classified into three different types based on their responses to depolarization current injections (Chieng et al., 2006; Dumont et al., 2002; Lopez de Armentia and Sah, 2004). The neurons that show a significant delay before firing action potentials are classified as late firing neurons (Fig. 3B) while neurons that fire without this delay are classified as regular spiking neurons (Fig. 3C). A third type of neurons do not display such delay but show bursting activity and sometimes a rebound firing post hyperpolarization are classified as low threshold bursting neurons. Confirmed previous findings, we found that late firing neurons and regular spiking neurons are mostly located in the lateral part of CeA while low-threshold bursting neurons are usually observed in the medial part of CeA. Because both the PKC-δ+ neurons and Htr2a+ neurons are located in the lateral part of CeA, we focus on the late firing neurons and regular spiking neurons. Comparing EPSC properties between these two populations, we did not observe any significant difference in EPSC amplitude or latency (Fig. 3J, K). Consistent with previous reports, the electrophysiological properties of these two types of neurons are significantly different (Fig. 3L-N). Together, these results suggest that the projection from the insular cortex to the CeA neurons is non-selective, and at the overall population level, insular neurons innervate all types of CeA neurons with similar strength despite their distinct genetic markers or electrophysiological properties.

### Activation of insula-CeA projections trigger action potential firing in CeA neurons

We next tested whether stimulating the insula-CeA projections differentially activates distinct populations of CeA neurons. When we current-clamped the CeA neurons and light-activated the ChR2-expressing terminals projected from insular cortex, we found that, in many cases, a brief light pulse can trigger action potential firing in CeA neurons (Fig. 4A). Although the EPSC amplitude is similar between the PKC-δ+ neurons and PKC-δ-neurons, fewer PKC-δ+ neurons (4 out of 11 neurons) than PKC-δ-neurons (6 out of 8 neurons) fired action potentials when the ChR2-expressing terminals were light-stimulated (Fig. 4B). If we consider only cells where action potentials were triggered, the successful action potential trigger rate is similar between these two populations (Fig. 4C). Interestingly, the PKC-δ-neurons show an adaptation in action potential triggering when stimulated at 10 Hz while PKC-δ+ neurons do not (Fig. S2). We then plotted the success rate of action potential triggering against the EPSC amplitude of the neurons. We found that a larger amplitude EPSC is required to trigger action potentials in CeA PKC-δ+ neurons than PKC-δ-neurons (Fig. 4D). Similar results were also observed in late firing neurons and regular spiking neurons (Fig. 4E-G). The different action potential triggering rates are consistent with their electrophysiological properties, i.e. the PKC-δ+ neurons or late firing neurons have a longer firing delay and therefore are more difficult to be triggered to fire action potentials. However, these results do not support the idea that CeA PKC-δ+ neurons are more likely to be activated when the insula-CeA pathway is light-stimulated and cannot explain why activation of the pathway suppresses food intake.

**Fig. 4.**
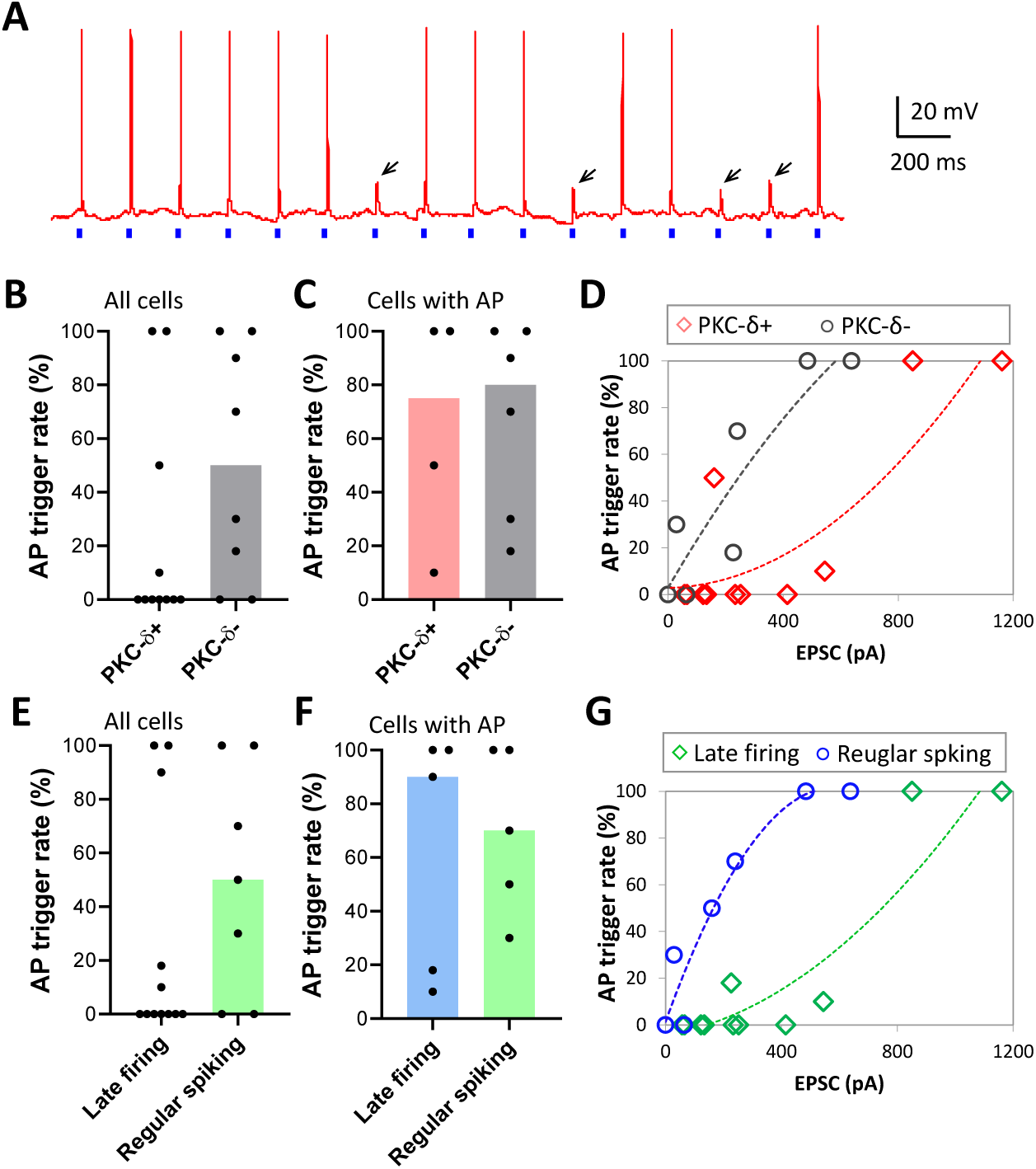
Activation of insular terminals in CeA triggers action potential firing in CeA neurons. A. Sample trace of current-clamp recording shows that action potential (AP) firing is triggered by 2-ms light pulses (blue dots). Note some light pulses cannot trigger action potential but a subthreshold depolarizing potential (indicated by arrows). B-C. Successful AP trigger rate in all the neurons (B) or neurons that at least one action potential is triggered (C). Bar graphs show median value. Each dot represents one cell. D. AP trigger rate is plotted against EPSC amplitude in PKC-δ+ neurons and PKC-δ-neurons. The dotted lines are polynomial fitting. Bar graphs show median value. Each data point represents one cell. E-F. Successful AP trigger rate in all the neurons (E) or neurons that at least one action potential is triggered (F). Bar graphs show median value. Each dot represents one cell. G. AP firing rate is plotted against EPSC amplitude in late-firing neurons and regular spiking neurons. The dotted lines are polynomial fitting. Bar graphs show median value. Each data point represents one cell.

### Using mathematical models to explore the functional connectivity structure of the CeA circuit

Experimental findings to be included in the model:

#### Finding 1

The PKC-δ+ and Htr2a+ neural populations in mice have similar electrophysiological properties though they play opposite roles in feeding. The majority of these cells are late firing neurons in both populations (Cai et al., 2014; Douglass et al., 2017).

#### Finding 2

Activating only the PKC-δ+ population leads to feeding suppression, while silencing this population leads to feeding promotion in a fed state (Cai et al., 2014).

#### Finding 3

Activating only the Htr2a+ population, which is a subset of the PKC-δ-population, leads to feeding promotion, while silencing this population (or silencing the PKC-δ-population) leads to feeding suppression (Cai et al., 2014; Douglass et al., 2017).

#### Finding 4

Surprisingly, activating both the PKC-δ+ and PKC-δ-population, which includes Htr2a+ neurons, through insular terminals consistently leads to feeding suppression (Fig. 1E, F).

The first three findings imply intuitively the following functional structure of the CeA circuit: the PKC-δ+ and Htr2a+ populations act in functionally opposite ways; PKC-δ+ leads to feeding suppression when activated, whereas Htr2a+ leads to feeding promotion when activated. However, given that these two populations are functionally opposing each other and share similar electrophysiological properties (Cai et al., 2014; Douglass et al., 2017; Haubensak et al., 2010), it is not clear how this functional structure can produce the outcome observed in Finding 4, in which the simultaneous activation of both the PKC-δ+ and Htr2a+ populations leads consistently to a winner-take-all situation where PKC-δ+ is *always* the dominating population, resulting in a net effect of feeding suppression. It is known that winner-take-all can occur through symmetry-breaking pitchfork bifurcation (Curtu et al., 2008; Werner and Spence, 1984); if so, the initial condition (and noise) would play a critical role in determining which of the two neural populations becomes the dominating one. A pair of identical and mutually inhibitory neuronal populations cannot *consistently* produce an outcome in which one particular population is always dominating the other as observed in Finding 4 (Rowat and Selverston, 1997; Shpiro et al., 2007). If we relax the assumption that these two populations are identical, such as by allowing for a sufficient amount of asymmetry between the two populations, then the winner-take-all outcome of Finding 4 can occur consistently (more on this later). However, existing experimental evidence does not suggest such strong asymmetry between the LF1 and LF2 populations (Cai et al., 2014; Douglass et al., 2017; Haubensak et al., 2010). A question arises: is it possible that a third population of neurons consisting of regular spiking neurons, most of which are negative for either PKC-δ or Htr2a (Cai et al., 2014; Douglass et al., 2017), plays a functionally significant role as part of the overall CeA circuit in modulating feeding behavior?

To reconcile the existing results and to compare various possible CeA circuit structures, we consider a simple neuronal network model of the CeA circuit consisting of three distinct CeA neurons: a late firing neuron (LF1) representing the PKC-δ+ neural population, another late firing neuron (LF2) representing either the Htr2a+ population or a subset of the PKC-δ-population, and possibly a third neuron that is regular spiking (RS) as another subset of the PKC-δ-population. Existing findings suggest that synaptic connections within the CeA circuit are inhibitory (Haubensak et al., 2010; Hou et al., 2016; Hunt et al., 2017; Sun and Cassell, 1993) and that the LF1 PKC-δ+ and LF2 PKC-δ-/Htr2a+ neurons are mutually inhibitory (Cai et al., 2014; Douglass et al., 2017; Haubensak et al., 2010). Hence, in this study we only allow for variations in the inhibitory synaptic connections from the third RS neuron to the two LF neurons as shown in Fig. 5A. Given the above assumptions, we need to consider four network connectivity scenarios shown in Fig. 5B, C, D, and E. Note that Fig. 5B is the special case in which the RS neuron is absent. To explore the functional significance of the regular spiking or late firing properties of the individual CeA neurons, we also consider variations in the electrophysiological properties of the three CeA neurons, such as the third neuron being late firing instead of regular spiking, as shown in Fig. 5C’, D’, and E’, or the scenarios in which the LF1 (representing PKC-δ+) and LF2 (representing PKC-δ-/Htr2a+) neurons are regular spiking instead of late firing, as shown in Fig. 5D^1^, D^2^, and D^3^. Analyzing these alternatives allows us to address whether the late firing property of the PKC-δ+ and Htr2a+ populations, and possibly the presence of a third regular spiking subpopulation, is a necessary condition for the observed CeA circuit output behaviors.

**Fig. 5.**
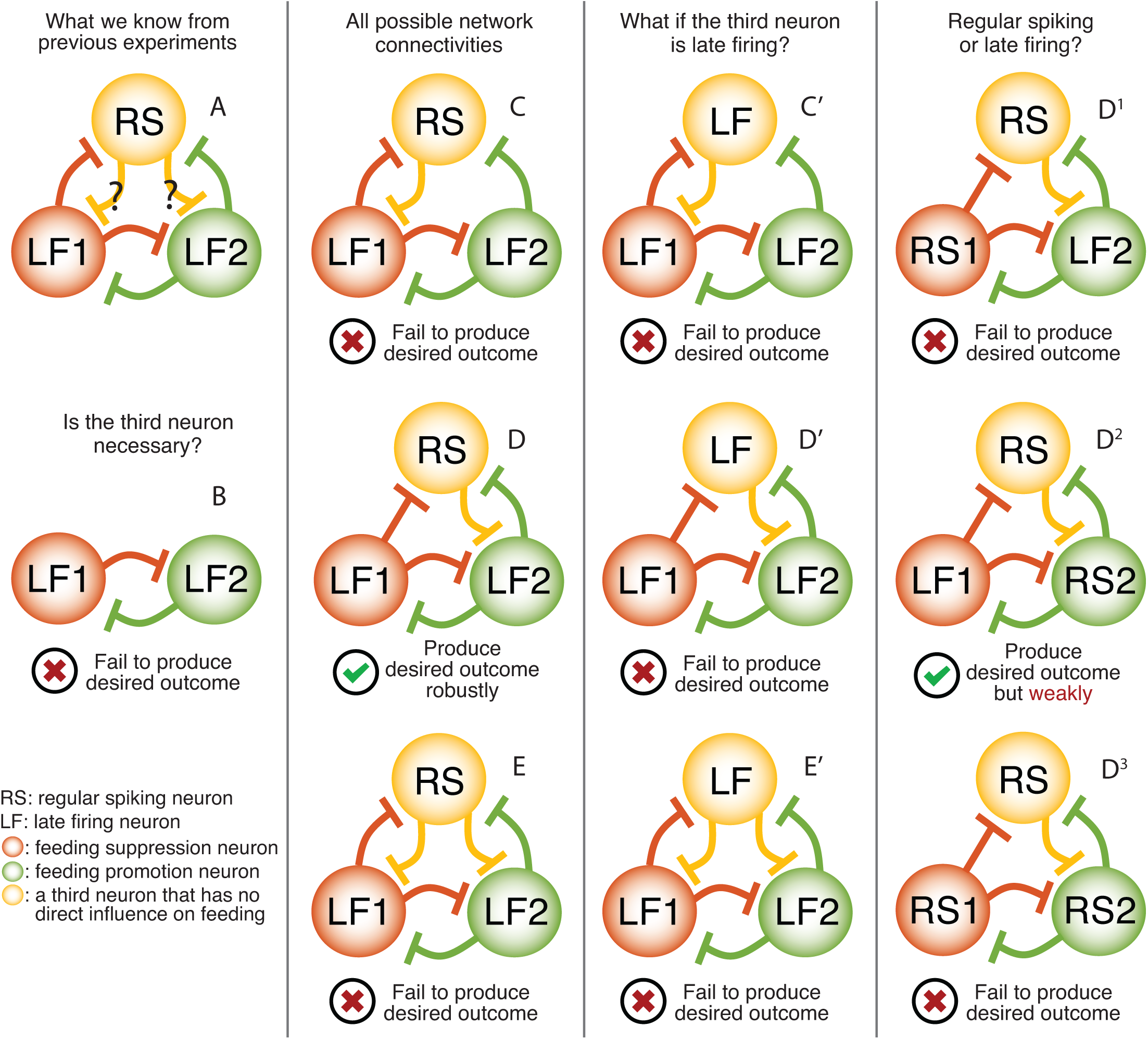
Various hypothetical circuit structures for the CeA circuit. We consider various combinations of connectivity topologies and intrinsic firing properties for the CeA circuit consisting of three CeA model neurons. Based on the model assumptions discussed in the Results section, symmetry analysis implies only nine fundamentally different scenarios (including the degenerated case with only two neurons, in which the third CeA neuron with no direct influence on feeding is absent). Computer simulations of our mathematical model of this CeA circuit shows that only the CeA circuit structure proposed in subfigure A is able to reproduce all previously observed feeding behavior change in response to various experimental conditions (see Fig. 6 for the computer simulation results of this scenario). Parameter values used in the computer simulations: 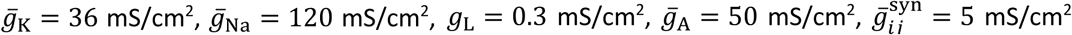 for all neurons, *E*_K_ = −77 mV, *E*_Na_ = 50 mV, *E*_L_ = −54.4 mV, *E*_syn_ = −70 mV, *τ*_*B*_ = 16 msec, *I*_bias_ = 20 mA, 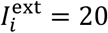 for activation, 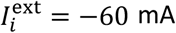 for silencing, 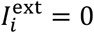 for the control condition, and *C* = 1 μF/cm^2^. Each simulation is run for 4000 msec; we only use results from the last one third of the time span to allow sufficient time for the stable activity to emerge from the initial transients.

Based on the CeA circuit structure proposed above, we develop a dynamical systems model of neuronal activity in the CeA circuit in which each of the three CeA neurons is represented by a Hodgkin-Huxley-type model neuron that produces the desired electrophysiological property (i.e., late firing or regular spiking) in each scenario (see Methods). We use an idealized conductance-based current model for the inhibitory synaptic connections with time-dependent channel dynamics mimicking the general behavior of GABA channels. Each model neuron receives an external input, which represents the activation input from the insular cortex; the external input is positive when the activation input is on and negative when the neuron is silenced. The resulting mathematical model of the entire CeA circuit is a system of differential equations with about 19 dynamic variables (the exact number of variables in each scenario varies and depends on the exact combination of the electrophysiological properties of the individual CeA neurons and the overall network connectivity scenario). Note that we do not consider the effect of heterogeneity within each CeA neural population in this article. In future work, we shall consider the heterogeneous case.

Finally, we describe how the CeA circuit output is linked to feeding behavior in our model. Findings 2 and 3 suggest the following simple model on the effect of the CeA circuit output on feeding behavior: the firing rate of LF1 neuron (representing the PKC-δ+ population) is proportional to the strength of feeding suppression, and the firing rate of LF2 neuron (representing the Htr2a+ population) is proportional to the strength of feeding promotion. We assume that the activity of RS neuron does not have a direct influence on feeding behavior. Consistent with our principle of using the fewest assumptions possible, we assume that the negative effect on feeding exerted by LF1 and the positive effect by LF2 sum up linearly, and this sum provides a quantitative measure of the net effect of the CeA circuit output on feeding.

### A third regular spiking subpopulation of neurons are functionally significant for producing the desired CeA circuit dynamics

Among the four scenarios shown in Fig. 5B, C, D, and E, we found only one specific CeA circuit structure capable of reproducing all four experimental findings. Fig. 5D shows this successful CeA circuit structure, in which the RS neuron receives an inhibitory synaptic connection from LF1 and forms a mutually inhibitory synaptic connection with LF2, and therefore providing a disynaptic inhibitory pathway from LF1 to LF2 in addition to the existing mutual inhibition between the LF1 and LF2 neurons. The second column in Fig. 6 shows the membrane potential time course of our three CeA neurons, using this CeA circuit structure, under various conditions. In the control condition (no additional external input to any of the three CeA neurons except for the baseline bias input), the extra disynaptic inhibitory pathway from LF1 via RS to LF2 breaks an otherwise perfect symmetry between LF1 and LF2, resulting in a firing pattern of 2:1—2:1—1:1 with LF1 firing more frequently than LF2. Intuitively, as we activate both LF1 and LF2 (by increasing the external drive to LF1 and LF2 but not to RS), the firing rates of both LF1 and LF2 become elevated, resulting in a net increase in the number of spikes fired by LF1 (since LF1 already fires more spikes than LF2 in the control condition). This net increase is further exacerbated by a change in the firing pattern of LF1 and LF2 from 2:1—2:1—1:1 to a persistent 2:1 ratio. Either of these two effects alone leads to a net increase in LF1 spikes, and therefore, activating both the LF1 and LF2 neurons results in feeding suppression. This result is shown in the second row in Fig. 6. The third row of Fig. 6 shows that the activation of LF1 leads to feeding suppression; in this case, an elevated LF1 neuron leads to an increased inhibition to LF2, resulting in a 3:1 ratio in the firing rate of LF1 over LF2. The fourth row in Fig. 6 shows the opposite case in which LF1 is silenced; in this case, the LF2 and RS neurons become the only active neurons, resulting in feeding promotion. The fifth row reproduces the result that LF2 activation leads to feeding promotion; the elevated activity of LF2 leads to a reversal in the firing rate ratio of LF1 over LF2 from the 2:1—2:1—1:1 pattern (control condition) to the 1:2—1:2—1:1 pattern, resulting in a net increase in the number of spikes of LF2, and therefore producing a net effect of feeding promotion. In the last row, as we silence only the LF2 neuron, LF1 and RS become the only active neurons, resulting in feeding suppression.

**Fig. 6.**
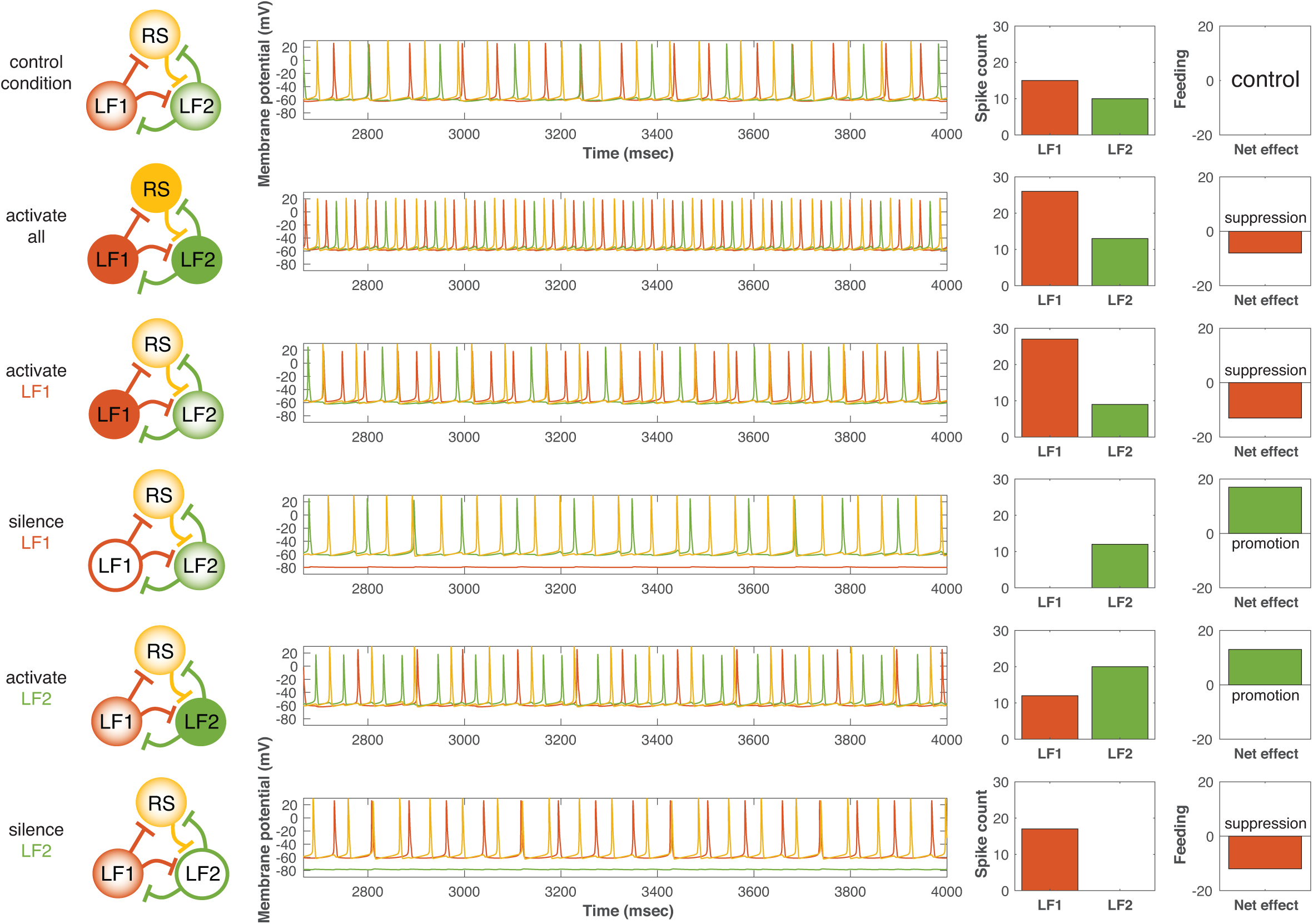
The response of our proposed CeA circuit model to various external stimuli. We subject our proposed CeA circuit model (shown in Fig. 5A) to five different conditions: activating all three model neurons 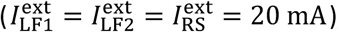, activating only the LF1 neuron 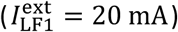, silencing only the LF1 neuron 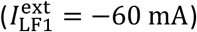, activating only the LF2 neuron 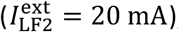, and silencing only the LF2 neuron 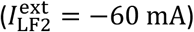. The first column shows the five different conditions (in addition to the control condition shown in the first row) imposed on our CeA circuit model. Each circle represents a CeA model neuron; a solid color circle indicates activation, a hollow color circle indicates silencing, and a color circle with a color gradient indicates the control condition 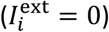. The second column shows the membrane potentials of the three CeA model neurons as functions of time; the red curve for the LF1 neuron, the green curve for the LF2 neuron, and the yellow curve for the RS neuron. Each simulation is run for 1000 msec. Note that the figure shows only the neuronal activities during the last one third of the time span to allow sufficient time for the initial unstable transient activity to settle to a stable activity. The third column shows the spike count of the LF1 and LF2 neurons (during the last one third of the time span). In the last column, we show the net effective of the CeA circuit firing activity on feeding behavior. Recall that in our model the firing rate of LF1 neuron is proportional to its effect on feeding suppression, and the firing rate of LF2 neuron is proportional to its effect on feeding promotion, whereas the dynamics of the RS neuron has no direct effect on feeding behavior (the RS neuron indirectly influence the feeding behavior through its synaptic connections with the other two neurons). The net effect is obtained by subtracting the LF1 neuron spike count from the LF1 neuron spike count (we treat the control condition as the neutral case; the net spike count in the control condition provides the baseline for comparison). All model parameter values are the same as described in Fig. 5 caption (except we use a longer time span here for illustration purposes).

We then consider several alternative synaptic organizations. We first ask what will happen if the regular spiking neural population is absent. In this case, the original three-cell CeA circuit is reduced to a pair of mutually inhibitory late firing neurons as shown in Fig. 5B. This symmetric mutual inhibitory structure gives rise to identical firing rate in both the LF1 and LF2 neurons. Not surprisingly, this scenario is able to reproduce experimental Findings 2 and 3, in which LF1 or LF2 is either activated or silenced. However, when both LF1 and LF2 are activated, the net effect on feeding is nil, since the elevated suppressing effect of LF1 is cancelled out perfectly by an equally elevated promoting effect of LF2. This outcome is a consequence of our model assumption that the LF1 and LF2 neurons are identical and that the two neurons are coupled through mutual inhibitory with equal strength. However, if we allow for a sufficient amount of asymmetry between LF1 and LF2, it is possible that the winner-take-all outcome observed in Finding 4 can occur. To see how much asymmetry is needed to reproduce Finding 4, we introduce two types of asymmetry between LF1 and LF2: (1) an asymmetry in the strength of the activation input to each neuron, and (2) an asymmetry in the mutual inhibition between the two neurons. Results are shown in Fig. 7. We find that, in order to produce all of the desired outcomes, including the winner-take-all outcome required by Finding 4, either the maximum conductance of the synaptic inhibition from LF1 to LF2 has to be at least twice of that from LF2 to LF1, or the external current input into LF1 has to be at least twice of that into LF2. This result suggests that in order for a two-cell (both late firing) network to produce all the desired firing-rate behaviors, a strong asymmetry between the LF1 and LF2 populations is needed. However, existing experimental evidence does not suggest such strong asymmetry between the LF1 and LF2 populations (Cai et al., 2014; Douglass et al., 2017; Haubensak et al., 2010).

**Fig. 7.**
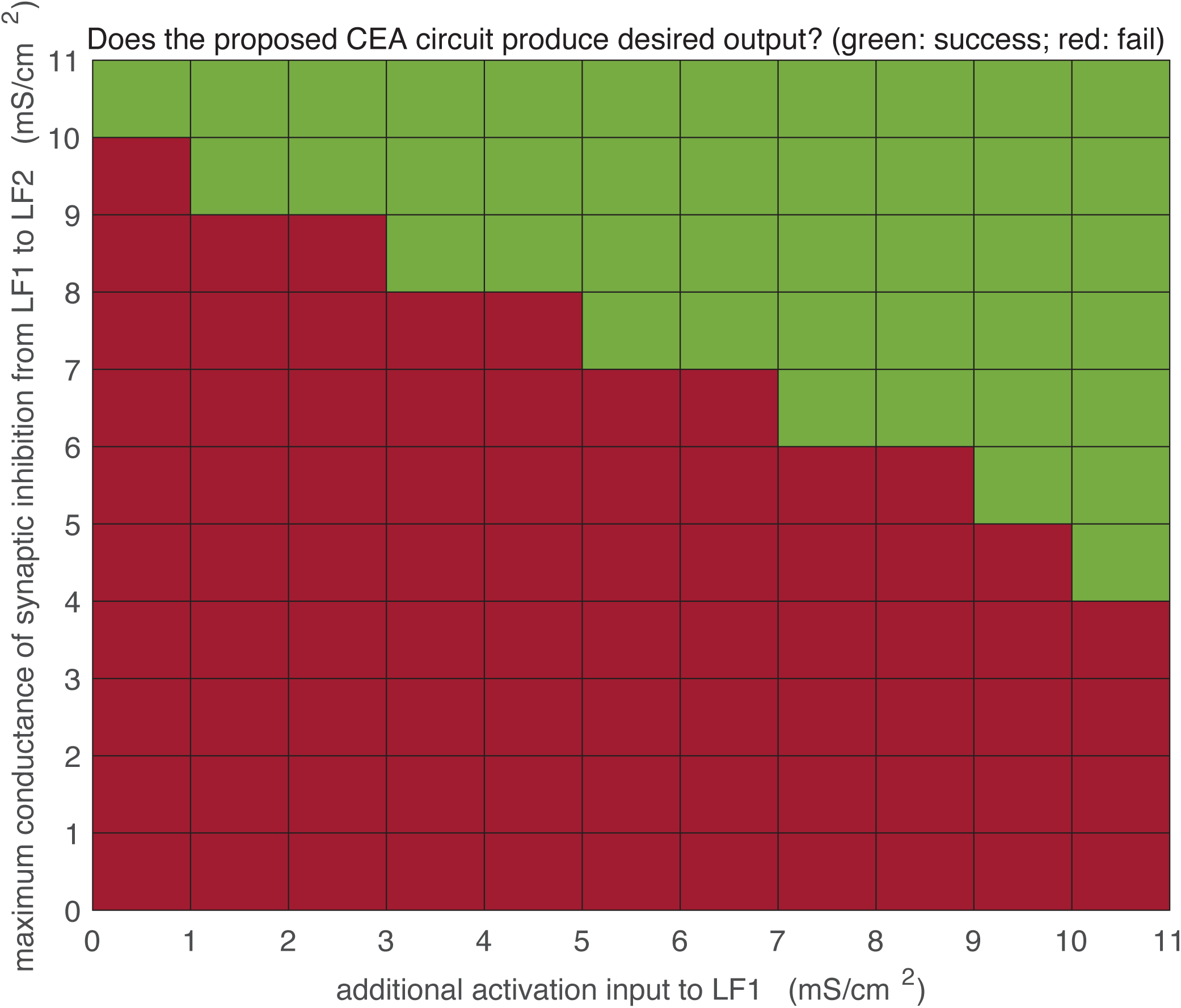
Effect of asymmetry in the two-cell LF1-LF2 network. Computer simulations of the two-cell network (LF1-LF2) with varying degrees of asymmetry. We introduce asymmetry to the two-cell network by adjusting two parameters: (1) the maximum conductance of the synaptic inhibition from LF1 to LF2 (introducing an asymmetry in the mutual inhibition between LF1 and LF2), and (2) an extra external current input to the LF1 neuron (introducing an asymmetry in the insular cortex inputs to LF1 and LF2). For each combination of parameter values, we run a series of computer simulations under the same five experimental conditions (as in Fig. 6) to test whether the CeA circuit being tested can reproduce all desired CeA circuit output outcomes. If successful (i.e., all desired outcomes are reproduced), a green block is plotted at the location corresponding to the parameter combination, and if unsuccessful (i.e., at least one desired outcome is not reproducible), a red block is plotted. All other parameters values are the same as described in Figs. 5 and 6 captions. In all computer simulations, in order to declare an outcome as feeding suppression, we require that the number of LF1 spikes to be at least three more than the number of LF2 spikes, whereas to declare an outcome as feeding promotion, we require that the number of LF2 spikes to be at least three more than the number of LF1 spikes.

We continue to consider two additional alternative connectivity scenarios. In Fig. 5C, the RS neuron is bidirectionally coupled to LF1 via mutual inhibition; in Fig. 5E, we consider a fully coupled network of LF1, LF2, and RS neurons, in which each neuron forms a mutual inhibition with the other two neurons. Computer simulations show that none of the above alternative scenarios is able to reproduce all of the existing experimental findings. Specifically, all of the alternative scenarios shown in Fig. 5B, C, and E can reproduce Findings 2 and 3 but fail to reproduce Finding 4. This suggests that the third regular spiking neuronal population plays a functionally significant role in modulating CeA circuit output, and that this modulation is in the form of an inhibitory pathway from the PKC-δ+ population to the regular spiking population and finally to the Htr2a+ population.

### The late firing property of the CeA neurons is necessary to produce the experimentally observed behaviors

We have identified a specific CeA circuit structure that reproduce qualitatively all of the four experimental findings. In particular, our proposed CeA circuit structure (Fig. 5D) consists of three neural populations, in which a mutually inhibitory pair of late firing neurons represents the PKC-δ+ population (LF1) and the Htr2a+ population (LF2), respectively, and a third regular spiking neural population inserts an additional inhibitory pathway from LF1 to LF2. Our model is consistent with the experimental finding that both the PKC-δ+ and Htr2a+ populations are mostly late firing neurons (Douglass et al., 2017; Haubensak et al., 2010). This raises another interesting question: is the late firing property of CeA neurons (represented by LF1 and LF2 neurons in our model) a necessary electrophysiological condition for the CeA circuit to produce the experimentally observed behaviors under various experimental conditions?

To address this question, we use our mathematical model to further explore various alternative scenarios by modifying the prescribed electrophysiological property of the PKC-δ+ and Htr2a+ populations in our model: in Fig. 5D^1^, we consider the case in which the PKC-δ+ neural population is regular spiking (instead of late firing); in Fig. 5D^2^, we consider the case in which the Htr2a+ population is regular spiking (instead of late firing); and in Fig. 5D^3^, we consider the case that both the PKC-δ+ and Htr2a+ neurons are regular spiking. Computer simulations show that scenarios D^1^ and D^3^ fail to reproduce Finding 4 (that the activation of both PKC-δ+ and PKC-δ-neurons should lead to feeding suppression), whereas scenario D^2^ is able to reproduce the desired outcome but in a weak manner (in the sense that the feeding suppression effect when both the PKC-δ+ and Htr2a+ populations are activated is much weaker than the suppression effect under the condition when only the PKC-δ+ population is activated). This result implies that the late firing property of the PKC-δ+ and Htr2a+ neurons is essential to ensure the desired feeding behavior. Note that a key feature of our model is that the regular spiking neuron does not have a direct influence on feeding behavior, however, the regular spiking neuron *does* have an indirect effect on the neuronal activity of the CeA circuit its effect on other populations, and therefore our result also suggests that the regular spiking property of the third neural population is also necessary to achieve the desired CeA circuit output behaviors.

To test the robustness of our modeling results against variations in the strength of synaptic connections between the CeA neurons, we performed a two-dimensional numerical bifurcation study to examine how the solutions of our dynamical systems model for the CeA circuit change as we vary the values of the parameters 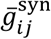, the strength of the synaptic connection from neuron *j* to neuron *i*. Specifically, we vary the values of two types of synaptic connections, those originating from the late firing neurons, and those originating from the regular spiking neuron. We reran our computer simulations for various combinations of these two synaptic strength values and tested each case to verify whether all desired outcomes are satisfied. Fig. 8A and B show the bifurcation study results for our proposed CeA circuit (as shown in Fig. 5D) and the weak scenario as shown Fig. 5D^2^. A green color block indicates that all desired findings can be reproduced under this specific parameter combination, whereas a red color black indicates at least one of the desired outcomes is violated. Our result shows that our proposed CeA circuit is able to reproduce all desired outcomes over a wide range of synaptic strength combinations, whereas the weak scenario has a smaller area of green blocks. Overall, this bifurcation study shows that our main conclusions from the mathematical model is robust against variations in synaptic strength.

**Fig. 8.**
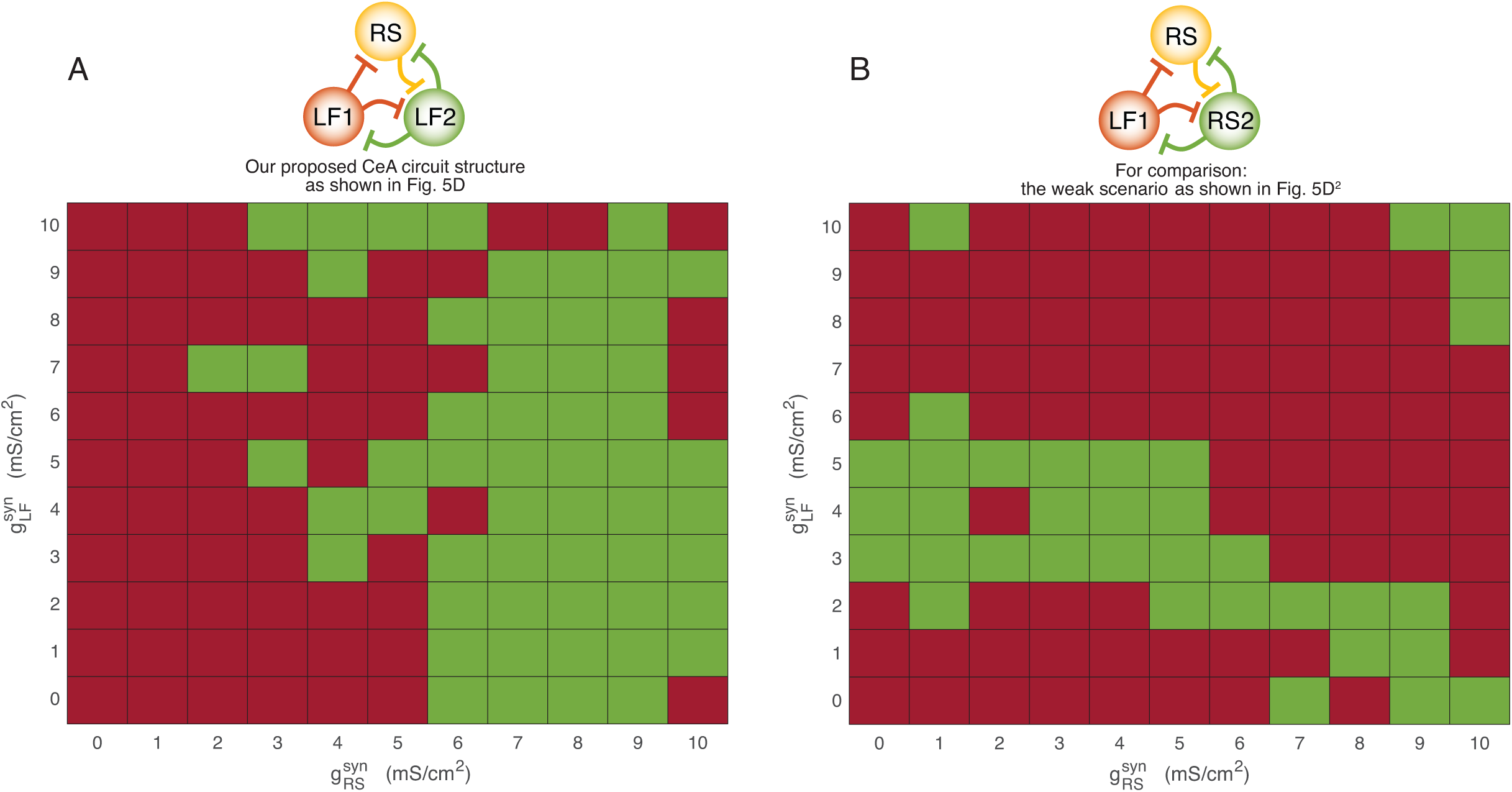
Robustness of our mathematical model results against variations in synaptic strength. For each combination of synaptic strength values, we run a series of computer simulations under the same five experimental conditions (as in Fig. 6) to test whether the CeA circuit being tested can reproduce all desired CeA circuit output outcomes. Using the same notation and criteria as described in Fig. 7 caption, if successful, a green block is plotted at the location corresponding to the parameter combination, and if unsuccessful, a red block is plotted. For comparison, we perform this two-dimensional numerical bifurcation study for two CeA circuit scenarios: in subfigure A, we test the scenario shown in Fig. 5D (i.e., our proposed CeA circuit structure); in subfigure B, we test the scenario shown in Fig. 5D^2^ (i.e., the scenario that produces only a weak outcome of feeding suppression when both the PKC-δ+ and Htr2a+ populations are activated). The parameter 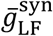 is the catch-all variable for all maximum conductance parameters 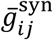 in which the presynaptic neuron *j* is late firing (regardless of the postsynaptic neuron *i*), whereas 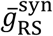 is the catch-all variable for all 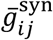 in which the presynaptic neuron *j* is regular spiking (regardless of the postsynaptic neuron *i*). All other parameters values are the same as described in Figs. 5 and 6 captions.

## Discussion

Neurons in CeA have been widely reported to play important roles in fear conditioning, anxiety, pain, reward processing, and feeding behaviors (Duvarci et al., 2011; Fadok et al., 2018; Gilpin et al., 2015; Janak and Tye, 2015; Johansen et al., 2011; LeDoux, 2000; Ressler and Maren, 2019; Thompson and Neugebauer, 2017). Accordingly, it is not surprising that many different types of CeA neurons were identified based on their distinct electrophysiological properties, neuropeptides or other genetic markers (Day et al., 1999; Dumont et al., 2002; Haubensak et al., 2010; Kim et al., 2017; van den Burg and Stoop, 2019). Recent studies using genetic marker-labelled CeA neurons revealed several subpopulations of CeA neurons with different functions in feeding regulation (Cai et al., 2014; Douglass et al., 2017; Hardaway et al., 2019; Ip et al., 2019; Kim et al., 2017). Especially, two non-overlapping CeA neurons, marked by the expression of PKC-δ+ and Htr2a+ respectively, were identified to regulate feeding in opposing direction (Cai et al., 2014; Douglass et al., 2017). Activation of CeA PKC-δ+ neurons suppresses food intake while activation of CeA Htr2a+ neurons increases food intake. Activation of CeA PKC-δ-neurons does not seem to affect food intake significantly, despite the Htr2a+ population being included in this subset (Cai et al., 2014; Douglass et al., 2017). Silencing CeA PKC-δ+ neurons increases food intake but silencing CeA PKC-δ-neurons or silencing CeA Htr2a+ neurons decreases food intake (Cai et al., 2014; Douglass et al., 2017). Surprisingly, a majority of the CeA PKC-δ+ neurons (86%) or Htr2a+ neurons (85%) in mice are late firing neurons (Cai et al., 2014; Douglass et al., 2017), suggesting that the distinct functions of these two populations are not purely a result of their electrophysiological properties. Neurons in the CeA receive inputs from diverse brain regions, some of which have been demonstrated to regulate behaviors related to feeding (Campos et al., 2016; Palmiter, 2018; Schiff et al., 2018; Wang et al., 2018). For example, it was reported that neurons in the lateral parabrachial nucleus (LPB) that express calcitonin gene-related protein (CGRP) relay danger information and project to the CeA to suppress food intake (Campos et al., 2018; Carter et al., 2013); neurons in insular cortex that encode bitter taste project to CeA to convey negative valence (Schiff et al., 2018; Wang et al., 2018). More recently, Gehrlach et al. expressed ChR2 in insular cortex under *CamK2a* promoter and demonstrated that activation of the insula→CeA pathway suppresses feeding, increases anxiety and induces other aversive behaviors (Gehrlach et al., 2019).Interestingly, when we used AAVretro-Cre in CeA and Cre-dependent ChR2 in insular cortex, we only observed feeding suppression but not anxiety or other aversive behaviors. This might reflect a weaker activation of insular cortex neurons, either a smaller number of neurons or a lower expression of ChR2 in the insular cortex could achieve a manipulation more specific to a certain behavior. This possibility was also supported by Gehrlach’s finding that a higher frequency (20 Hz) stimulation triggers immobility and many aversive behaviors while a lower frequency (10 Hz) does not (Gehrlach et al., 2019). Stern et al. recently found that inhibition of the projection from insular cortex Nos1 neurons to CeA has no effect on normal feeding, but blocks the context conditioned overconsumption, however, the connections from Nos1 neurons to different CeA neurons were not directly mapped (Stern et al., 2019). Retrograde rabies tracing studies showed that both the CeA PKC-δ+ and CeA PKC-δ-neurons receive monosynaptic inputs from many common brain regions including insular cortex and LPB (Cai et al., 2014; Douglass et al., 2017). Here our results demonstrate that neurons from insular cortex send monosynaptic excitatory inputs to all the CeA neurons non-selectively. The synaptic strength is not significantly different between CeA PKC-δ+ neurons and CeA PKC-δ-neurons, or between late firing neurons and regular spiking neurons. It seems that the CeA PKC-δ+ neurons require a larger EPSC to activate them than the CeA PKC-δ-neurons. Thus, it is surprising that activation of the neural pathway from insular cortex to CeA suppresses food intake, an effect similar to the activation of CeA PKC-δ+ neurons or silencing CeA PKC-δ-neurons. Therefore, the different functions of CeA neurons cannot be explained by their inputs either. Instead, these results suggest that the distinct functions of the CeA neurons might due to their special circuit structure within CeA.

The functional connections among the CeA neurons have been recently studied using two different strategies. One is based on the ChR2-assited circuit mapping, in which ChR2 is expressed in a subtype of CeA neurons with specific genetic markers and postsynaptic responses of other CeA neurons are recorded in response to light stimulation of the ChR2-expressing neurons. It was found that almost all the CeA PKC-δ-neurons receive monosynaptic inhibition from CeA PKC-δ+ neurons (Haubensak et al., 2010) and almost all CeA Htr2a negative (CeA Htr2a-) neurons are inhibited by light activation of the CeA Htr2a+ neurons (Douglass et al., 2017). Similarly, almost all CeA somatostatin (SOM) negative (CeA SOM-) neurons are inhibited by CeA SOM+ neurons (Hunt et al., 2017; Li et al., 2013). These studies indicate that CeA neurons form extensive inhibition with each other at the population level. The other strategy is based on paired whole-cell recording and stimulation on individual neurons, which also showed extensive interconnections among all electrophysiological types of CeA neurons (Hou et al., 2016; Hunt et al., 2017). Both unidirectional and bidirectional connections between late firing neurons, from late firing to regular spiking neurons, and from regular spiking to late firing neurons were observed (Hunt et al., 2017; Li et al., 2013). However, current available genetic markers cannot label all the different types of CeA neurons. For example, what genetic marker may label regular spike neurons is still unknown and the role of regular spiking neurons in feeding remains to be determined. The paired recording cannot detect the connections between cells that are far away from each other or cells located in different brain slices, and the number of cells that can be sampled with this method is usually limited. Therefore, it is difficult to measure the exact connections among different types of CeA neurons that have distinct functions with current technologies.

Although current experiments cannot tease out the exact CeA circuit connectivity, mathematical modeling is powerful in testing all possible network structures based on current knowledge of CeA neurons (functions, electrophysiological properties, and connections) to explore which specific neuronal network can produce the experimentally observed results. Constrained by existing experimental findings, we have built a conductance-based model for each type of CeA neurons using Hodgkin-Huxley-type equations, and used it to explore how the firing properties of individual CeA neurons and the CeA circuit’s overall synaptic organization combine to produce the desired circuit output for feeding control. Computer simulations of our mathematical model show that, in order to produce the experimentally observed feeding behaviors in response to various conditions, the presence of both late firing and regular spiking neurons are necessary. Furthermore, by examining different combinations of circuit connectivity scenarios, we find one specific CeA circuit synaptic organization that can reproduce previously known experimental findings. An important outcome of our CeA circuit model is that, at the control condition, the LF1 neuron (representing the PKC-δ+ population) has a higher firing rate than the LF2 neuron (representing the PKC-δ- or Htr2a+ population) (Ciocchi et al., 2010), so that the activation of both CeA populations (such as by the activation of insular cortex neurons) would lead to a net increase in the spikes fired by the LF1 neurons, which leads to feeding suppression according to our model assumption.

Note that, however, our proposed three-neuron structure is not necessarily the only possible structure to achieve the desired outcome. The necessity of a third regular spiking neuron is a consequence of our model assumption that the two late-firing LF1 and LF2 neurons are identical and that these two neurons are coupled through mutual inhibitory with equal strength. This result shows the potential functional significance of having a mixture of late firing and regular spiking neurons.

We would like to point out some limitations of our mathematical model. First, our CeA circuit model consists of three Hodgkin-Huxley-type model neurons, each of which represents a CeA neural population. Therefore, our model is not a population model and cannot not address heterogeneity within each CeA neural population or subregions of CeA or even outputs of the CeA neurons; in future work, we shall consider population models, such as Wilson-Cowan-type models (Destexhe and Sejnowski, 2009; Wilson and Cowan, 1973) or models based on large dynamical systems (Rangan and Young, 2013) that can take into account the heterogeneity within each CeA neural population. Second, the delayed onset firing property of our late firing CeA model neuron is implemented by the incorporation of a potassium A-type current, which is a fast-activation and slow-inactivation outward current (Rush and Rinzel, 1995). Other dynamical mechanisms, such as those caused by a slow calcium buildup (Rush and Rinzel, 1995; Terman et al., 2002), can also lead to a delayed onset firing, which we did not consider in our model. Third, our model does not consider long-term adaptations in the synaptic connections except our model includes time-dependent synaptic dynamics to account for the general behavior of GABA channels. It is possible that synaptic strength adapts to feeding behavior through certain feedback pathways, which requires further experiments to determine. Fourth, our model assumes that the firing rate of the LF1 neuron is linearly proportional to the strength of feeding suppression, and the activity of LF2 is linearly proportional to the strength of feeding promotion, and that these two effects add up linearly to give the net effect of the CeA circuit on feeding. We do not consider gain control mechanism or nonlinearity in our model. We did so because we aimed to use the fewest assumptions that are consistent with the existing experimental observations. The main feature of our CeA circuit model is that the feeding suppressing effect of the LF1 neuron directly competes with the feeding promoting effect of the LF2 neuron, and this competition is modulated by the mutual inhibition between these two late firing neurons, and at the same time, by the extra inhibitory pathway from the LF1 neuron through the RS neuron to the LF2 neuron.

While the actual connections within the CeA are still unknown, using computational modeling, we found that a specific circuit structure, combined with the intrinsic electrophysiological properties of the CeA neurons, can explain the puzzling feeding-related results. This circuit diagram can provide a platform for identifying specific types of CeA neurons involved in regulating feeding and other behaviors such as fear conditioning (Haubensak et al., 2010; Isosaka et al., 2015). From a broader perspective, a large amount of modeling work has studied the dynamics of individual conductance-based model neurons and the networks formed by these neurons (Izhikevich, 2007), such as in the stomatogastric ganglion (Selverston, 2008), the crayfish swimmeret system (Skinner and Mulloney, 1998; Zhang et al., 2014), and the hippocampus (Rich et al., 2016). However, to the authors’ knowledge, no previous study has examined the dynamics of a neural circuit consisting of at least two mutually inhibitory neurons in which one is regular spiking and the other is late firing. Hence, it is possible that the circuit structure and the underlying mechanism identified herein may be more broadly applicable.

## STAR+METHODS

Detailed methods are provided in the online version of this paper and include the following:

- Key resources table
- Lead contact and materials availability
- Method details
  - Mice
  - Virus
  - Stereotaxic mice surgery
  - In vivo Optogenetics
  - Feeding assays
  - Open field test
  - Histology
  - Electrophysiological slice recordings
  - Quantification and statistical analysis
  - Mathematical model

## Acknowledgements

We thank W. Haubensak and D. Anderson for PKC-δ-Cre mice; K. Deisseroth, E. Boyden for viruses; C. Fang for managing mice colony and genotyping; K. Gothard, S. Cowen for comments and critical reading of the manuscript. Part of the brain slice electrophysiology experiments were carried out in the laboratory of Dr. D. Anderson (California Institute of Technology). HC is supported by a NARSAD Young Investigator Grant from the Brain & Behavior Research Foundation, a grant from the Foundation for Prader-Willi Research and a grant from The Klarman Family Foundation Eating Disorders Research Grants Program.

## Author Contributions

CZM performed mathematical modeling and computer simulations, MS performed behavioral experiments, and HC performed electrophysiology experiments. CZM, MS, and HC wrote the paper.

## STAR Methods

### Key resources table

| REAGENT | SOURCE | IDENTIFIER |
| --- | --- | --- |
| Virus |  |  |
| AAV2-EF1a-DIO-hChr2(H134R)-EYFP-WPRE-pA | UNC, Deisseroth |  |
| AAV2-EF1a-DIO-EYFP-WPRE-pA | UNC, Deisseroth |  |
| AAV2-CMV-Cre-WPRE-SV40 | UNC, Deisseroth |  |
| AAV2-hSyn-ChR2-EYFP | UNC, Deisseroth |  |
| rAAV2-Retro/CAG-Cre | UNC, Boyden |  |
| Organisms/Strains |  | Stock # |
| Wild-type mice (C57BL/6) | Charles River Laboratories | C57BL/6 |
| PKC-δ-ires-Cre line | Caltech, David Anderson lab | |
| Ai 14 Cre reporter mice | Jackson Lab | 007914 |

### Contact for reagent and resource sharing

Requests for further information or biological resources and reagents should be directed to and will be fulfilled by Haijiang Cai. Request for mathematical modeling and Matlab coding should be directed to and will be fulfilled by Calvin Zhang-Molina.

### Mice

All mice used in this project are offspring from PKC-δ-Cre mice crossed with wild type C57BL/6crl mice from the Charles River Laboratory (a background used in our previous study (Cai et al., 2014)) for at least 5 to 6 generations. The genotype of offspring was identified by PCR on genomic tail DNA. Only male wild type and PKC-δ-Cre offspring were used in this study. All mice were housed on a 12-hour light (7 am)/dark (7 pm) cycle with *ad libitum* access to water and rodent chow unless placed on a food restriction schedule for fasted feeding experiments. All behavioral experiments and brain slice electrophysiology were performed during the light cycle. All animal care and experimental procedures were strictly conducted according to the guidelines of US National Institutes of Health for animal research and were approved by the Institutional Animal Care and Use Committee (IACUC) at the University of Arizona.

### Virus

All the viruses used in this study were purchased from the University of North Carolina Viral Vector Core. AAV2retro-Cre vector was originally generated in Dr. Ed Boyden’s lab at M.I.T. AAV2-EF1a-DIO-EYFP-WPRE-pA, AAV2-EF1a-DIO-hChR2(H134R)-EYFP-WPRE-pA, AAV2-hSyn-ChR2-EYFP, AAV2-CMV-Cre-WPRE-SV40 vectors were generated in Dr. Karl Deisseroth’s lab at Stanford University. All the AAV and AAV2retro viruses had titres of 1-6×10^12^ genome copies per ml. Construct validity and correct targeting to the brain nucleus of interest were confirmed through post-mortem processing of brain sections in multiple sets of mice or electrophysiological recordings on live brain slices. To minimize variation of environmental differences and virus expression level, the control virus and experimental virus were injected in the same time window by the same investigator.

### Stereotaxic mice surgery

The surgery procedure was described previously (Cai et al., 2014). In brief, mice at 2-3 months old were deeply anaesthetized with 5% isoflurane in oxygen and kept at 1-1.5% isoflurane throughout the surgery. Survival surgery was then performed on a stereotaxic frame (Model 1900, Kopf Instruments). An incision was made on the midline of the scalp and a craniotomy was performed above the target regions. Viruses were microinfused through a pulled-glass micropipette with 20-50 µm tip outer diameter connected with a Nanoliter Injector (Nanoliter 2010, World Precision Instruments) at a rate of 8-10 nl/min. After injection, the micropipette was raised by 50 µm and left in place for 3-5 min to allow for diffusion of the virus liquid before the pipette was slowly withdrawn. Injection volumes into the IC and CeA were 40 nl. Viruses were injected bilaterally for behavioral studies and unilaterally for slice electrophysiology. Injection coordinates (in mm) relative to midline and Bregma: insular cortex (±3.90, 0, −3.83), CeA (±2.85, −1.40, −4.73). Optical ferrule fibers were implanted bilaterally ∼0.5 mm above the injection coordinates. After ferrule fiber implantation, dental cement (C&B Metabond) was used to secure the fiber to the skull. For postoperative care, mice were injected intraperitoneally with ketoprofen (5 mg/kg) daily for 3 days. At least 3 weeks after surgery were allowed for recovery and viral expression before the behavioral assays.

### *In vivo* Optogenetics

A blue (Shanghai DreamLaser: 473 nm, 100 mW or 50 mW) or yellow (Shanghai DreamLaser: 593 nm, 100 mW) laser was used to deliver light stimulation. An Accupulser Signal Generator (World Precision Instruments, SYS-A310) was used to control the frequency and pulse width of the laser light. Light was delivered to the brain through an optic fiber (200 µm diameter, NA 0.22, Doric Lenses) connected with the implanted ferrule fiber by a zirconium sleeve. The light power in the brain regions 0.5 mm below the fiber tip was calibrated as previously described (Aravanis et al., 2007). The calibrated light power density (0.5 mm below the fiber tip) used in light activation experiment was ∼5 mW/mm^2^. 10 Hz, 10-ms (pulse width) light pulse trains were used in optogenetic activation experiments.

### Feeding assays

Mice were transferred into an empty testing cage in the behavioral testing room to habituate for at least 20 min one day before the feeding test. For the 24-hr fasted feeding test, mice were food-deprived while have access to water. Mice were briefly anaesthetized with isoflurane (< 1 min) and coupled with optic fibers before the experiments. At least 20 min after recovery in behavioral testing room, mice were introduced into a clean empty testing cage with a pre-weighed regular food pellet, and allowed to feed for 20 min. The body weight of the mice before test, weight of food pellet before and after test, including the food debris left in the cage floor after test, were measured to calculate the amount of food intake. For the feeding test at fed state, mice were not food deprived before testing, and allowed to feed for 30 min. For optogenetic experiments, the light was delivered just after the mice were introduced into the testing cage. After each test, mice were returned to their home cage with *ad libitum* access to water and rodent chow. For the home cage feeding test, mice were food deprived for 24 hours. A single food pellet was placed in the home cage at the beginning of the test and the animal was allowed to eat for 10 min. Activation light (473 nm) was triggered 1-2 seconds after each feeding behavior began. 10 Hz, 10-ms light pulses were delivered until 1-5 seconds after the cessation of each feeding. The feeding behavior was videotaped and manually analyzed with a MATLAB based in-house behavioral annotation script.

### Open field test

A white square box (50 × 50 × 30 cm, a 25 × 25 cm square center was defined as “center” in analysis) was used as open field box. Mice were placed individually in the center of the box, and their behavior was tracked for 6 min in optogenetic tests with 2 minutes of light stimulation (473 nm, 10 ms pulse, 10 Hz) applied 2 minutes after the start. All the behaviors were videotaped and analyzed offline with Ethovision.

### Histology

All mice after behavioral tests were deeply anesthetized with ketamine/xylazine (100/20 mg/ml). Mice were then transcardially perfused with 20-ml PBS followed by 20-ml of 4% paraformaldehyde in PBS. Brains were removed and post-fixed in 4% paraformaldehyde overnight before being rinsed twice with PBS. The brains were sectioned with a vibratome (Leica, VT1000S) at 50-100 µm thickness and plated for imaging. The expression of virus and position of implanted optic fibers were checked with a fluorescence microscope.

### Electrophysiological slice recordings

Mouse brain slice electrophysiology experiments were performed as described previously (Cai et al., 2014). Brain coronal sections were sectioned at 250 µm thickness with a Leica vibratome (VT1000S) in ice cold glycerol-based artificial cerebrospinal fluid (GACSF) containing 252 mM glycerol, 1.6 mM KCl, 1.2 mM NaH_2_PO_4_, 1.2 mM MgCl_2_, 2.4 mM CaCl_2_, 18 mM NaHCO_3_, 11 mM glucose, oxygenated with carbogen (95% O_2_ balanced with CO_2_ for at least 15 min before use. The brain sections after cutting were recovered for at least one hour at 32-34 °C in regular ACSF containing 126 mM NaCl, 1.6 mM KCl, 1.2 mM NaH_2_PO_4_, 1.2 mM MgCl_2_, 2.4 mM CaCl_2_, 18 mM NaHCO_3_, 11 mM glucose, oxygenated with carbogen. The recordings were performed in a rig equipped with a fluorescence microscope (Olympus BX51), MultiClamp 700B and Digidata 1550A1 (Molecular Devices). The patch pipettes with a resistance of 5-10 MΩ were pulled with P-97 Sutter micropipetter puller and filled with an intracellular solution (135 mM potassium gluconate, 5 mM EGTA, 0.5 mM CaCl_2_, 2 mM MgCl_2_, 10 mM HEPES, 2 mM MgATP and 0.1 mM GTP, pH 7.3-7.4, 290-300 mOsm). Recording data were sampled at 10 kHz, filtered at 3 kHz and analyzed with pCLAMP10. tdTomato expression were usually verified post recording using a RFP filter. For the optogenetic stimulation, a blue laser (Shanghai DreamLaser, 473 nm, 50 mW) was used to deliver light pulses (0.1-2 mW/mm^2^ at the tip). 2 ms light pulses were used to activate the nerve terminals from insular cortex to trigger action potentials or induce postsynaptic responses in CeA neurons. EPSCs were measured when cells were voltage-clampped at −70 mV, a 0.1 µs 0.1 mV was applied at the same time of light delivery to help identify the start of light pulse. The action potentials were triggered when cells were current-clampped at −65 mV.

### Quantification and statistical analysis

Unless indicated, Data represent mean ± s.e.m. Unpaired Student’s *t*-test or Mann-Whitney rank test was used to compare two groups. A *p* value smaller than 0.05 was considered significant. Data were analyzed with GraphPad Prism Software, R and RStudio (538.1 Qt/5.4.1).

### Mathematical model

In this subsection, we introduce and describe our mathematical model of the CeA circuit. We first describe the mathematical equations for each model neuron in the CeA circuit. In our model, the state of each model neuron is completely described by its membrane potential and the gating variables of its intrinsic and synaptic ionic currents, whose activities are altogether governed by a set of Hodgkin-Huxley-type equations that produce a prescribed electrophysiological property of either regular spiking or late firing. Here, the phrase “late firing” refers to a relatively long *onset firing wait time*, which measures the elapsed time from when a depolarization current step is injected to the cell to when the cell fires an action potential (if the depolarization current was sufficient to trigger the firing event). Experimental results have shown that onset delay recordings from central amygdala neurons fall into two clusters: one with short delays (or almost no delay), and the other with long delays (Chieng et al., 2006; Dumont et al., 2002; Lopez de Armentia and Sah, 2004). Those central amygdala neurons that exhibit short or no delay in onset firing are called regular spiking CeA neurons, and those with a long onset delay are called late firing CeA neurons.

For each of the regular spiking CeA model neurons, we use the standard Hodgkin-Huxley equations (Ermentrout and Terman, 2010):

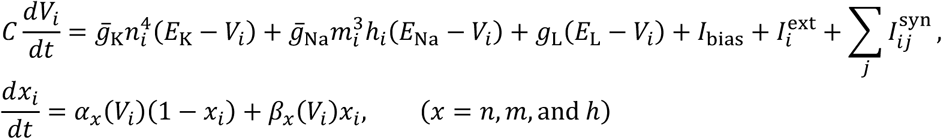

in which *V*_*i*_ is the membrane potential of model neuron *i* (*i* can be LF1, LF2, RS, LF, RS1, or RS2), *n*_*i*_ is the activation gating variable of the K^+^ current, *m*_*i*_ is the activation gating variable of the Na^+^ current, *h*_*i*_ is the inactivation gating variable of the Na^+^ current, 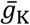 is the maximum conductance of the K^+^ current, 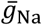 is the maximum conductance of the Na^+^ current, *g*_L_ is the conductance of the leakage current, 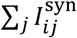 is the sum of all postsynaptic currents in neuron *i* due to activity in neuron *j* (*j* can be LF2, LF1, RS, LF, RS2, or RS1; this synaptic current will be defined later), *E*_K_, *E*_Na_ and *E*_L_ are the reversal potentials of the K^+^, Na^+^, and leakage currents, respectively, *I*_bias_ is the bias current, 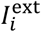 is the external current injection representing the activation input from the insular cortex to each neuron, *C* is the membrane capacitance, and the voltage-dependent gating functions are

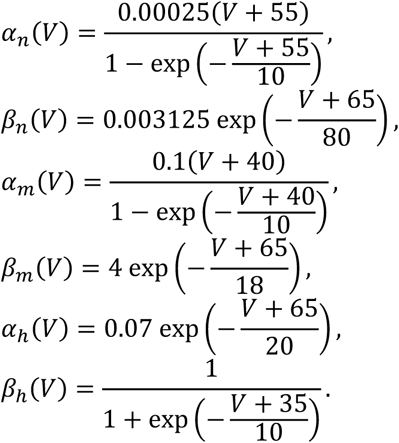

See figure captions for parameter values used in the computer simulations. Note that we use the same parameters for all computer simulations unless otherwise stated when performing a bifurcation analysis.

For each of the late firing CeA model neurons, we modify the standard Hodgkin-Huxley equations by including a potassium A-type current to the membrane potential equation (see the first term on the right-hand-side of the equation):

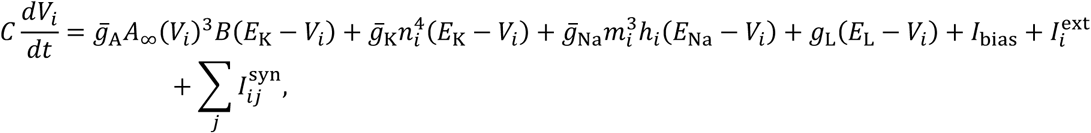

where the time evolution of the inactivation gating variable *B* of this A-type K^+^ current is governed by

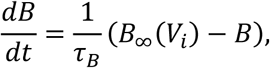

in which *τ*_*B*_ is the time constant for the inactivation of this A-type current, and the activation and inactivation gating functions are given by, respectively,

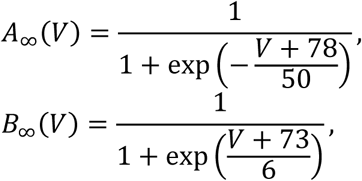

and all the other ionic currents are governed by the same equations as defined earlier in the case of a regular spiking neuron. Note that the inclusion of this fast-activation and slow-inactivation outward A-type current gives rise to a dynamical mechanism that produces delayed onset firing (Rush and Rinzel, 1995); by adjusting the time constant *τ*_*B*_ of the slow current inactivation, we can achieve a good range of spike onset delay (see Fig. 7).

Finally, we describe the mathematical equations for the synaptic connections among the CeA model neurons. We use a conductance-based synaptic current model for each inhibitory synaptic connection in which the gating variable itself is a dynamic variable following first-order kinetics (recall that experimental evidence suggests the lack of excitatory connections among the different CeA neural populations):

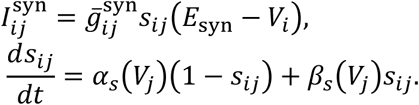

Recall that 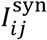 is the postsynaptic conductance in neuron *i* due to activity in neuron *j*, in which 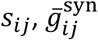 and *E*_syn_ are the gating variable, maximum conductance and reversal potential of this synaptic current, respectively, and the voltage-dependent gating functions are

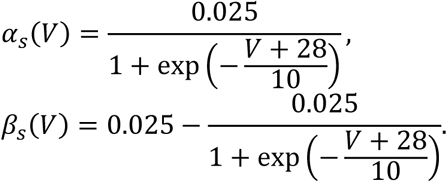

---

**Figure S1.**
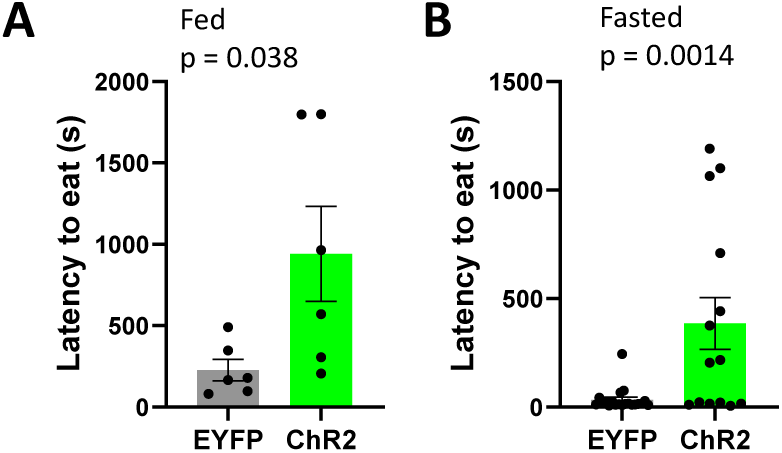
Initiation of feeding was suppressed in both fed (A) and fasted states (B). Light activation of the insula-CeA projections increases the latency to eat. The latency was defined from the presence of the food pellet to the first bite. Unpaired *t*-test, fed state, t_(10)_ = 2.39. n = 6 animals in each group; t_(32)_ = 3.50. n = 20 animals expressing EYFP and 14 animals expressing ChR2.

**Figure S2.**
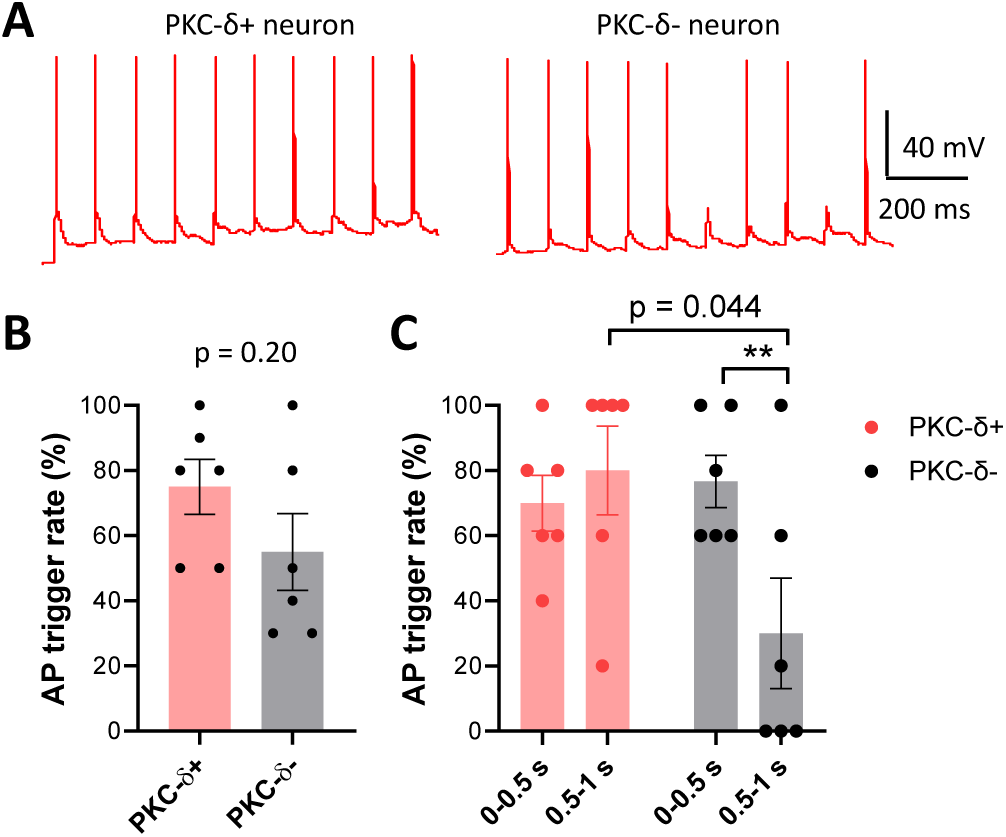
Action potentials (AP) in CeA neurons triggered by 10 Hz light stimulation of the insular terminals in CeA. **A**. Representative brain slice electrophysiological recordings show AP in PKC-δ+ and PKC-δ-neurons. **B**. The overall AP trigger rate is not different between PKC-δ+ and PKC-δ-neurons. Unpaired *t*-test, t_(10)_ = 1.38. n = 6 cells in each group. **C**. PKC-δ-neurons show a decreased AP trigger rate (adaption) while PKC-δ+ neurons does not. For PKC-δ-neurons, paired *t*-test, t_(5)_ = 4.18 ** p < 0.01, n = 6 cells in each group. Comparison at 0.5-1 s between PKC-δ+ and PKC-δ-neurons, unpaired *t*-test, t_(10)_ = 2.30. n = 6 cells in each group.

